# ASPECT: Alternative Splicing Event Classification with Transformers

**DOI:** 10.64898/2026.02.04.700904

**Authors:** Sahil Thapa, Miguelangel Tamargo, Oluwatosin Oluwadare

## Abstract

**Motivation:** Alternative splicing (AS) is a fundamental regulatory mechanism that expands transcriptomic and proteomic diversity by generating multiple mRNA isoforms from a single gene. Aberrant AS has been implicated in numerous diseases through the production of dysfunctional or pathogenic protein variants. However, much of the existing AS research has focused predominantly on exon skipping and constitutive splicing, with comparatively limited attention to other biologically relevant AS event types. Moreover, many current computational approaches rely on short genomic sequence windows and conventional deep learning architectures, which may limit their ability to capture broader regulatory context associated with complex splicing decisions. Bridging these methodological and conceptual gaps is essential for advancing comprehensive AS characterization and improving our understanding of its role in human health and disease.

**Results:** We present ASPECT, an alternative splicing event classification framework built upon DNABERT-2 with Byte Pair Encoding (BPE) tokenization. Across multiple binary alternative splicing event pair classification tasks, ASPECT achieves consistently strong performance as measured by AUC, F1-score,and accuracy, demonstrating reliable discrimination between closely related splicing event types. Importantly, ASPECT demonstrates consistent performance when applied to TCGA BRCA cancer-associated splicing events reconstructed from SpliceSeq annotations, supporting its applicability beyond the canonical splicing events used for training.

**Availability:** The open-source code, data, and detailed documentation used in this study are available at https://github.com/OluwadareLab/ASPECT.

**Supplementary information:** N/A

## 1 Introduction

Alternative splicing is a vital post-transcriptional process that increases proteomic and functional complexity in higher eukaryotes by allowing multiple mRNA and protein isoforms to be generated from a single gene [1] (Fig. 1). Precise classification of alternative splicing isoforms is important for understanding gene expression regulation and function. However, alternative splicing produces a vast complexity of isoforms that has posed challenges for comprehensive identification and classification. Early studies found evidence of alternative splicing but underestimated its widespread influence, with estimates that over 90% of human multi-exon genes undergo alternative splicing [2]. High-throughput sequencing techniques transformed the field by enabling genome-wide analyses of transcriptomes. These RNA-seq experiments revealed far greater alternative splicing complexity than anticipated, with thousands of genes expressing multiple isoforms in a tissue or condition-specific manner [3].

**Fig. 1:**
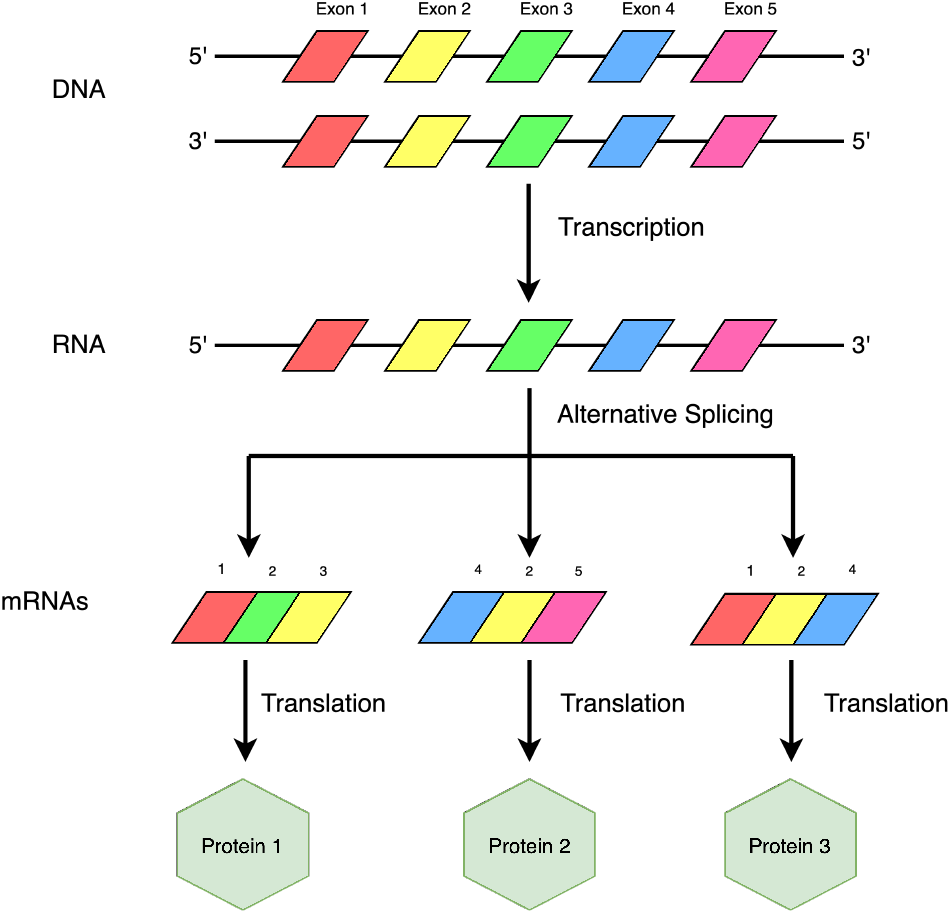
Overview of alternative splicing. DNA is transcribed into an RNA precursor, which can undergo alternative splicing to generate multiple mature mRNA isoforms that are subsequently translated into distinct protein products.

Alternative splicing introduces regulated variation into RNA processing by altering splice-site and exon–intron usage during pre-mRNA maturation [4] (Fig. 2). Several major classes of alternative splicing events contribute to transcript diversity, including *exon skipping* (also referred to as *cassette exon* events), in which an exon may be selectively included or omitted from the mature transcript, and *alternative* 5^*′*^ or 3^*′*^ *splice-site* selection, which modifies exon boundaries by shifting splice junctions [5]. In contrast, *constitutive splicing* represents the default splicing pattern in which exons are consistently retained, producing a canonical mRNA isoform. Through regulated alternative splicing, a single gene can generate multiple protein isoforms with distinct and sometimes opposing biological functions, substantially expanding proteomic complexity in eukaryotic systems.

**Fig. 2:**
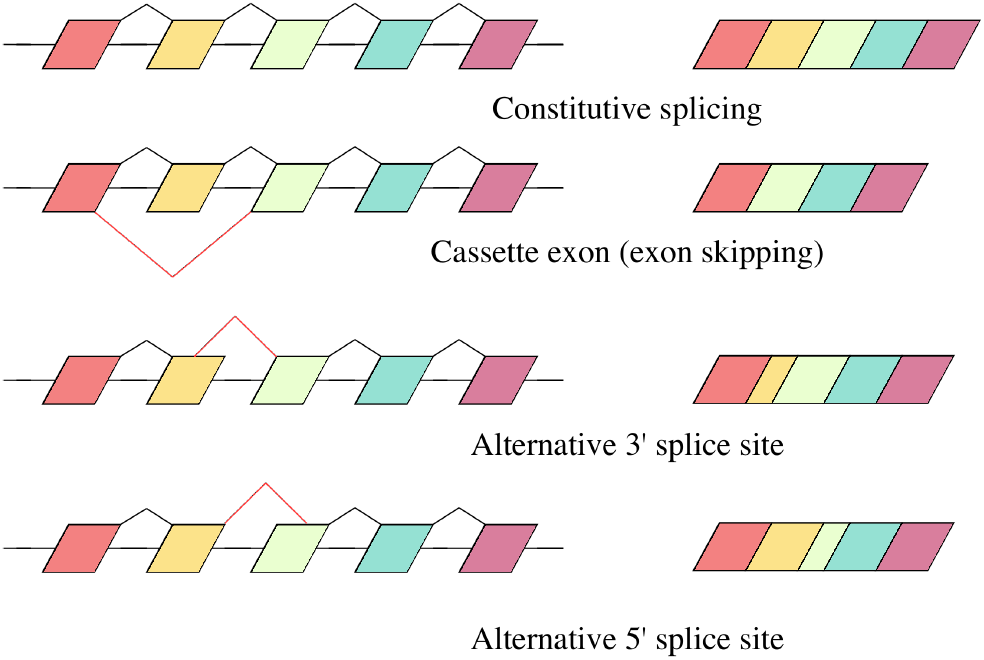
Schematic illustration of common splicing event types including constitutive splicing, cassette exon (exon skipping), alternative 3^*′*^ splice site, and alternative 5^*′*^ splice site.

Key factors shown to influence isoform characteristics include splice site strengths, exon/intron architecture, and phylogenetic conservation around exon boundaries. Integrative analyses have also linked splicing patterns to genomic features like cis-regulatory elements to provide mechanistic insights into splicing regulation [6]. Computational analysis of this massive volume of data presented new difficulties in reliably detecting and cataloging all isoforms. Advancements in bioinformatics tools have facilitated more sophisticated classification of alternative splicing patterns. Early tools focused on differentiating major annotated transcript variants but struggled with novel isoforms [7].

Existing approaches to alternative splicing (AS) classification can be broadly categorized into reference-free methods, that is, they do not rely on reference genome, and reference-based methods, they rely on reference genome for prediction. Reference-free approaches enable splicing classification in organisms or datasets lacking high-quality reference genome assemblies and include methods such as MCTASmRNA [8], DeepASmRNA [9], AStrap [10], and MkcDBGAS [11], which operate directly on transcript or sequence-derived representations. In contrast, reference-based algorithms, such as Deep Splicing Code [12] and DeepSplice [13], leverage reference genomes or aligned genomic context to inform splicing classification and to model underlying regulatory signals. **Although each category offers distinct strengths, reference-based approaches remain critical for accurately modeling complex splicing patterns in well-annotated genomes**. In particular, accurately resolving overlapping and closely related splicing event types continues to be challenging, motivating the development of more sophisticated reference-based models capable of capturing complex regulatory dependencies.

Existing reference-based computational approaches for alternative splicing (AS) event identification include the work of Busch and Hertel (2015) [14], which employed support vector machines (SVMs), as well as more recent deep learning frameworks such as the Deep Splicing Code (DSC) proposed by Louadi et al. (2019) [12] and DeepSplice proposed by Abrar et al. (2024) [13], both of which utilize convolutional neural networks (CNNs) for splicing event classification. Prior work has noted that distinguishing constitutively spliced exons from highly included alternative exons can be particularly challenging, as these events often exhibit similar local sequence patterns and splice-site strengths, motivating evaluation across exon inclusion or usage levels [12].

Despite substantial progress in computational modeling of alternative splicing, accurate classification of AS events remains an open problem. In particular, separating constitutive splicing from high-inclusion alternative splicing patterns is difficult because splicing regulation depends not only on core splicesite motifs but also on multiple cis-regulatory elements distributed across exons and flanking intronic regions. Consequently, models that rely primarily on locally constrained or fixed-length sequence representations may have limited capacity to capture broader contextual signals relevant to splicing regulation [3]. The importance of incorporating extended genomic context for splicing-related prediction tasks has been further demonstrated by approaches such as SpliceAI, which leverage long-range sequence information to improve splice-site prediction accuracy [15].

To address these limitations, we propose *Alternative Splicing Event Classification with Transformers* (ASPECT), a transformer-based framework built upon DNABERT-2 [16] with Byte Pair Encoding (BPE) tokenization. This design enables flexible and efficient representation of long nucleotide sequences, alleviating the rigidity and scalability constraints associated with fixed *k*-mer encoding schemes. By modeling fixed-length splice-junction–centered sequence windows of 1,024 base pairs, ASPECT incorporates both local splicing signals and extended genomic context, thereby improving discrimination across multiple alternative splicing event types.

## 2 Methods and Materials

### 2.1 Data and Preprocessing

The data curation strategy employed in this study follows the established framework proposed by Busch and Hertel [14] for constructing high confidence alternative splicing (AS) event datasets. To ensure that each extracted instance corresponds to a single, well defined splicing configuration, we applied additional quantitative filtering criteria based on expressed sequence tag (EST) evidence.

For exon skipping (cassette exon) events, exon utilization was quantified using the standard *inclusion level* metric [14]:

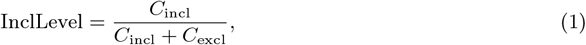

where *C*_incl_ and *C*_excl_ denote the number of ESTs supporting exon inclusion and exclusion, respectively.

For alternative 3^*′*^ and 5^*′*^ splice site events, splice junction preference was measured using the *usage level* :

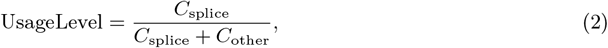

where *C*_splice_ represents the number of ESTs supporting the splice junction of interest and *C*_other_ corresponds to ESTs supporting alternative junctions.

Splicing event annotations were obtained from the HEXEvent database [17], which maps transcript isoforms to the hg38 human reference genome using curated UCSC Genome Database [18] annotations. HEXEvent additionally provides a strict filtering mode that retains only events supported by a unique EST configuration, thereby reducing transcript ambiguity and improving annotation reliability.

Based on these criteria, the final filtering strategy was defined as follows:

- **Constitutive splicing**: events with no supporting evidence for alternative splicing.
- **Cassette exon (exon skipping)**: events with an inclusion level below 90%.
- **Alternative 3**^*′*^ **splice sites**: events with `3usageLevel` values within the interval [0, 1].
- **Alternative 5**^*′*^ **splice sites**: events with `5usageLevel` values within the interval [0, 1].

Table 1 summarizes the distribution of splicing event classes following filtration.

**Table 1.** Distribution of splicing event classes after HEXEvent-based filtration.

| AS Event Type | Constitutive | Cassette | 3' AS | 5' AS |
| --- | --- | --- | --- | --- |
| Count | 41,204 | 12,874 | 2,629 | 1,331 |

Sequence preprocessing was performed using the human reference genome (`hg38.fa`) in conjunction with the `pybedtools` Python library to extract exon–intron junction sequences based on genomic coordinates [19]. All candidate events were first validated to ensure compliance with the inclusion and usage level criteria described above. Duplicate entries originating from the HEXEvent dataset were subsequently removed, and all nucleotide sequences were standardized to uppercase to ensure compatibility with the downstream tokenization and modeling pipeline. Each validated splicing event was then assigned to its corresponding splicing category (constitutive, cassette exon, alternative 3^*′*^, and alternative 5^*′*^) and stored in separate CSV files containing the fixed-length sequence (1,024 bp) and its associated event label. These curated datasets were used for train–validation–test partitioning and served as direct inputs to DNABERT-2 for tokenization and embedding during model fine-tuning.

### 2.2 ASPECT model Architecture

In this section, we describe the three major components of the ASPECT pipeline (Fig. 3).

**Fig. 3:**
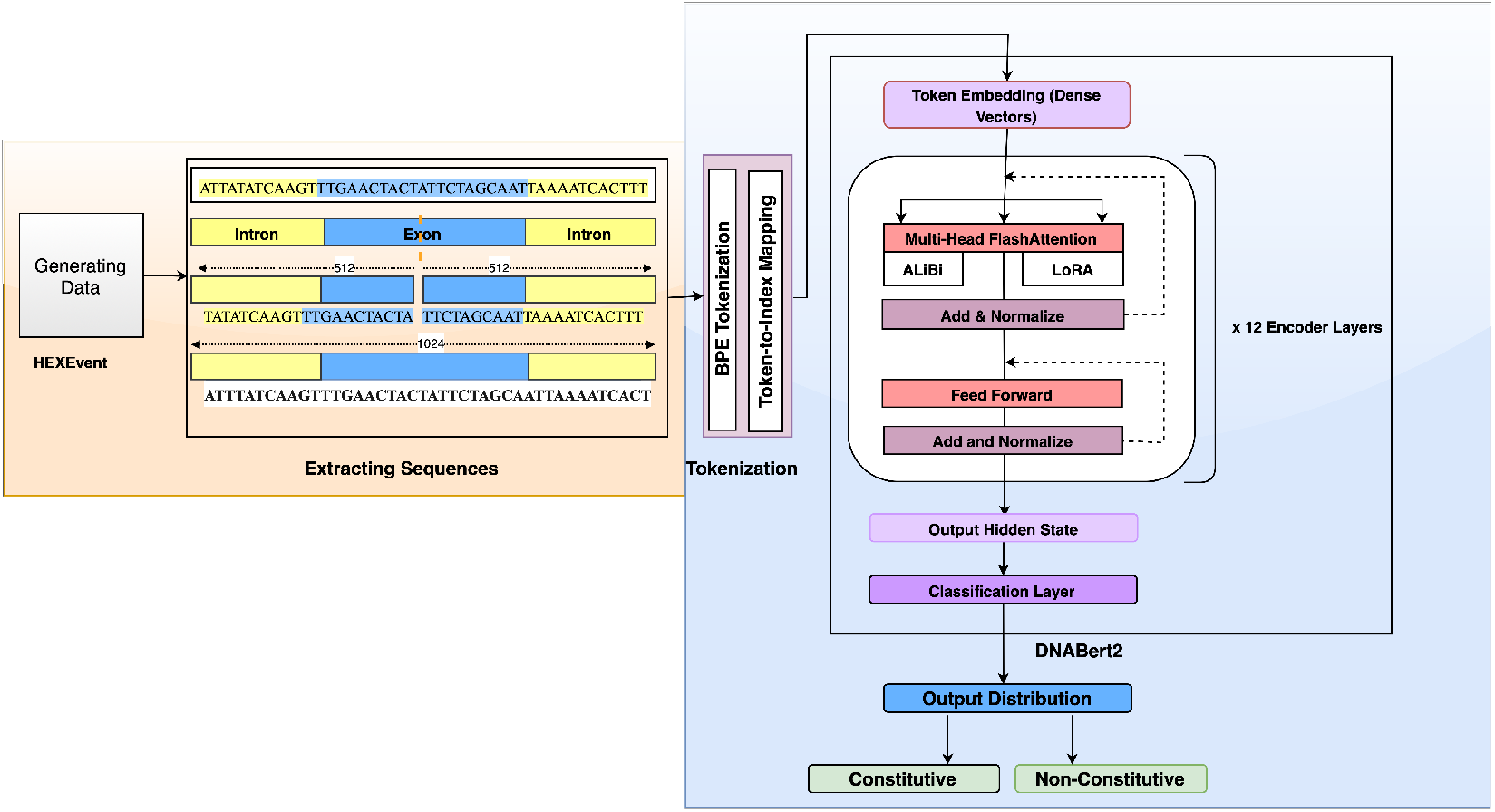
Overview of the ASPECT pipeline. Alternative splicing events curated from the HEXEvent database are converted into fixed length DNA sequence representations and tokenized using Byte Pair Encoding (BPE). The resulting token sequences are embedded with DNABERT-2 and processed by a multi-layer transformer encoder. A classification head produces posterior probabilities over splicing event classes.

#### Features Selection

To mitigate the impact of potential sequencing or annotation artifacts that may result in abnormally short exons or introns, we applied length-based filtering criteria, excluding regions shorter than 25 nucleotides for exons and 80 nucleotides for introns, following prior observations [20]. For each retained splicing event, we extracted fixed-length junction-centered sequence windows consisting of 512 nucleotides upstream and 512 nucleotides downstream of the exon. This procedure yields a single contiguous 1,024 base pair sequence per event, providing sufficient genomic context to capture both local splice site signals and longer range regulatory elements. While this study adopts a 1,024 bp window, the framework is readily extensible to alternative context lengths.

#### Input Encodings

##### BPE-based DNABERT-2

We adopt Byte Pair Encoding (BPE) tokenization through DNABERT-2[16] to represent genomic sequences in a flexible and scalable manner. BPE [21] performs data-driven subword segmentation by iteratively merging frequently occurring nucleotide patterns, enabling the model to capture both short regulatory motifs and longer contextual dependencies without requiring a predefined *k*-mer size. This approach substantially reduces input dimensionality and memory overhead while preserving biologically meaningful sequence structure, thereby improving the model’s capacity to learn hierarchical representations from DNA sequences.

#### Pre-Trained Transformer Model

DNABERT-2 builds upon the original DNABERT architecture [22] and supports alternative tokenization strategies, including BPE, while being pretrained on substantially larger and more diverse genomic corpora. It employs a 12-layer bidirectional Transformer encoder and is pretrained using a masked language modeling objective, enabling the model to learn contextualized representations of DNA sequences. Through self-attention, DNABERT-2 effectively captures long-range dependencies across intronic and exonic regions, which is critical for accurately modeling splicing regulatory elements such as splice sites, enhancers, and silencers. In our experiments, each input consists of a fixed-length nucleotide sequence of 1024 base pairs, providing sufficient genomic context to capture both local sequence motifs and distal regulatory signals relevant to alternative splicing. We initialize ASPECT using publicly available DNABERT-2 pretrained weights and finetune all model parameters on our filtered, labeled dataset to predict alternative splicing event classes.

### 2.3 Training, Validation, and Performance Evaluation

We employ a train–validation–test split strategy, rather than *k*-fold cross-validation, to streamline training and hyperparameter optimization. The dataset is partitioned into training (60%), validation (20%), and test (20%) subsets, with the validation set used exclusively for model selection and tuning.

During fine-tuning, we initialize the model with pretrained DNABERT-2 weights and optimize it using the Adam/AdamW optimizer with a cross-entropy-related objective. Hyperparameters, including learning rate, batch size, weight decay, number of training epochs, and loss-related parameters, are tuned using the validation set. These loss-related parameters are used to emphasize harder-to-classify splicing patterns and improve discrimination between closely related alternative splicing event types. DNABERT-2’s classification head outputs posterior probabilities over alternative splicing event classes.

Hyperparameter optimization is performed using the Optuna framework [23], which conducts an automated search over the defined parameter space using 18 independent trials. For each trial, a model is trained with a unique hyperparameter configuration and evaluated on the validation set, with the best-performing checkpoint selected based on the validation metric. This approach enables efficient exploration of the hyperparameter space while avoiding information leakage from the test set.

Model performance is assessed using standard classification metrics, including accuracy, F1-score, and area under the ROC curve (AUC), reported on test data. Confusion matrices are additionally used to analyze misclassification patterns, particularly between closely related splicing event classes.

All training experiments were conducted on a dedicated workstation equipped with two NVIDIA RTX 4090 GPUs (24 GB VRAM each), 125 GB of system memory, and a multi-core CPU, running Ubuntu 24.04.3 LTS.

### 2.4 Evaluation Metrics

We evaluate model performance using standard classification metrics, including F1-score, accuracy, and area under the receiver operating characteristic curve (AUC).

**F1-score** is defined as the harmonic mean of precision and recall:

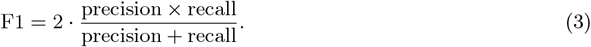

**Accuracy** measures the proportion of correctly classified samples:

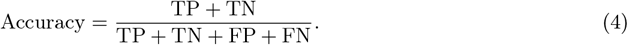

**AUC** summarizes the trade-off between the true positive rate (TPR) and false positive rate (FPR) across thresholds and is estimated as the area under the ROC curve:

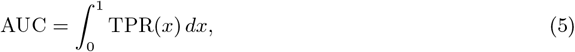

where *x* denotes the false positive rate.

## 3 Results

Model performance was evaluated using three classification metrics, Area Under the ROC Curve (AUC), F1-score, and Accuracy, reported separately for each binary classification task (Table 2). To ensure consistency across all experiments, fixed-length input sequences of 1,024 nucleotides were used during both training and testing.

**Table 2.** Performance of ASPECT across binary splicing event classification tasks, evaluated using AUC, F1-score, and Accuracy.

| Classification Task | AUC | F1-score | Accuracy |
| --- | --- | --- | --- |
| Constitutive vs. Cassette | 0.9526 | 0.9112 | 0.9114 |
| Constitutive vs. Alt 3' | 0.7732 | 0.7038 | 0.7072 |
| Constitutive vs. Alt 5' | 0.6725 | 0.6314 | 0.6315 |
| Cassette vs. Alt 5' | 0.8932 | 0.8495 | 0.8496 |
| Cassette vs. Alt 3' | 0.9037 | 0.8418 | 0.8422 |
| Alt 3' vs. Alt 5' | 0.6229 | 0.5880 | 0.5883 |

All auxiliary biological annotations, such as exon inclusion levels (e.g., high versus low), were intentionally excluded from the input feature space throughout the entire workflow. This design choice ensures that the model learns exclusively from intrinsic sequence-level information, thereby improving its robustness and generalizability to real world settings where such auxiliary labels are often unavailable.

To further characterize classification behavior beyond aggregate performance metrics, we analyzed confusion matrices for all binary classification tasks (Figure 4). Comparisons involving cassette exons show high true-positive rates and limited misclassification, whereas tasks distinguishing alternative splice-site events exhibit greater confusion. In particular, discrimination between alternative 3^*′*^ and alternative 5^*′*^ splice sites remains the most challenging, reflecting the intrinsic similarity of their underlying splicing signals (Table 2).

**Fig. 4:**
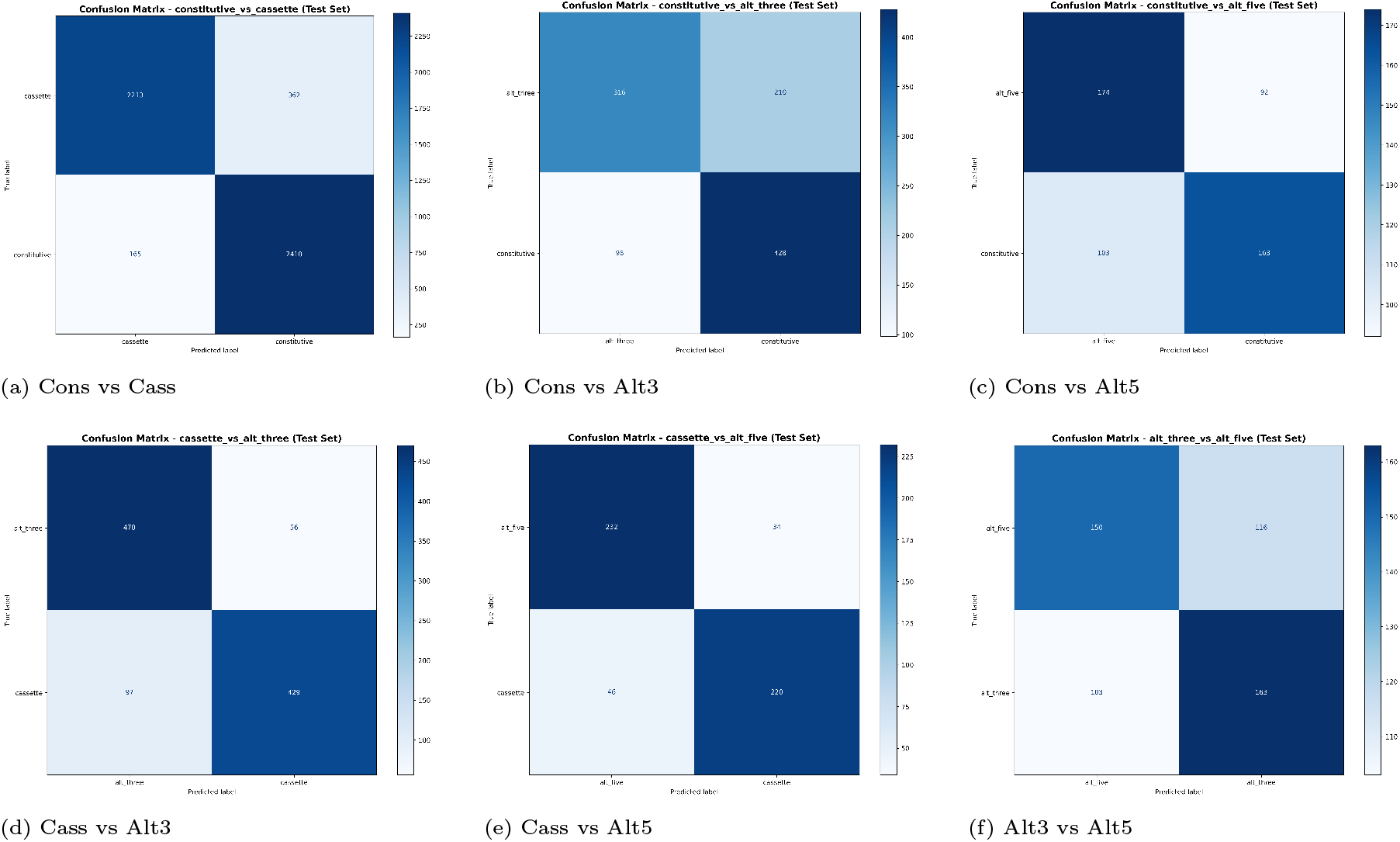
Confusion matrices for all six binary splicing event classification tasks evaluated using ASPECT on the test set. Comparisons involving cassette exons, including constitutive (Cons) versus cassette (Cass) (a), cassette versus alternative 3^*′*^ (Alt3) (d), and cassette versus alternative 5^*′*^ (Alt5) (e), exhibit strong separability with high true positive rates. Tasks distinguishing constitutive exons from alternative splice-site events show moderate performance, with constitutive versus Alt3 (b) performing consistently and constitutive versus Alt5 (c) exhibiting increased misclassification. The most challenging task is discriminating between Alt3 and Alt5 splice sites (f), reflecting the intrinsic similarity of their splicing signals.

### 3.1 Benchmarking with State-of-the-art Algorithm

To assess the performance of ASPECT, we benchmarked it against a recent state-of-the-art model for alternative splicing event prediction in the human genome, DeepSplice [13]. DeepSplice is designed to operate on short genomic windows (approximately 283 bp), focusing on local splice-site context, whereas ASPECT is trained to leverage substantially longer sequence contexts (1,024 bp) in order to capture both proximal and distal regulatory signals relevant to splicing. Accordingly, each model was evaluated using its native input sequence length during benchmarking, reflecting their intended operating conditions.

This evaluation setting enables a comparison of the practical performance of the two models as end-to-end alternative splicing classifiers, while acknowledging architectural differences in the amount of sequence context utilized. Figure 5 presents the receiver operating characteristic (ROC) curves for DeepSplice (orange) and ASPECT (blue) across multiple binary alternative splicing event pair classification tasks. ASPECT consistently achieves higher area under the curve (AUC) values, indicating stronger discriminative capability under their respective design settings.

**Fig. 5:**
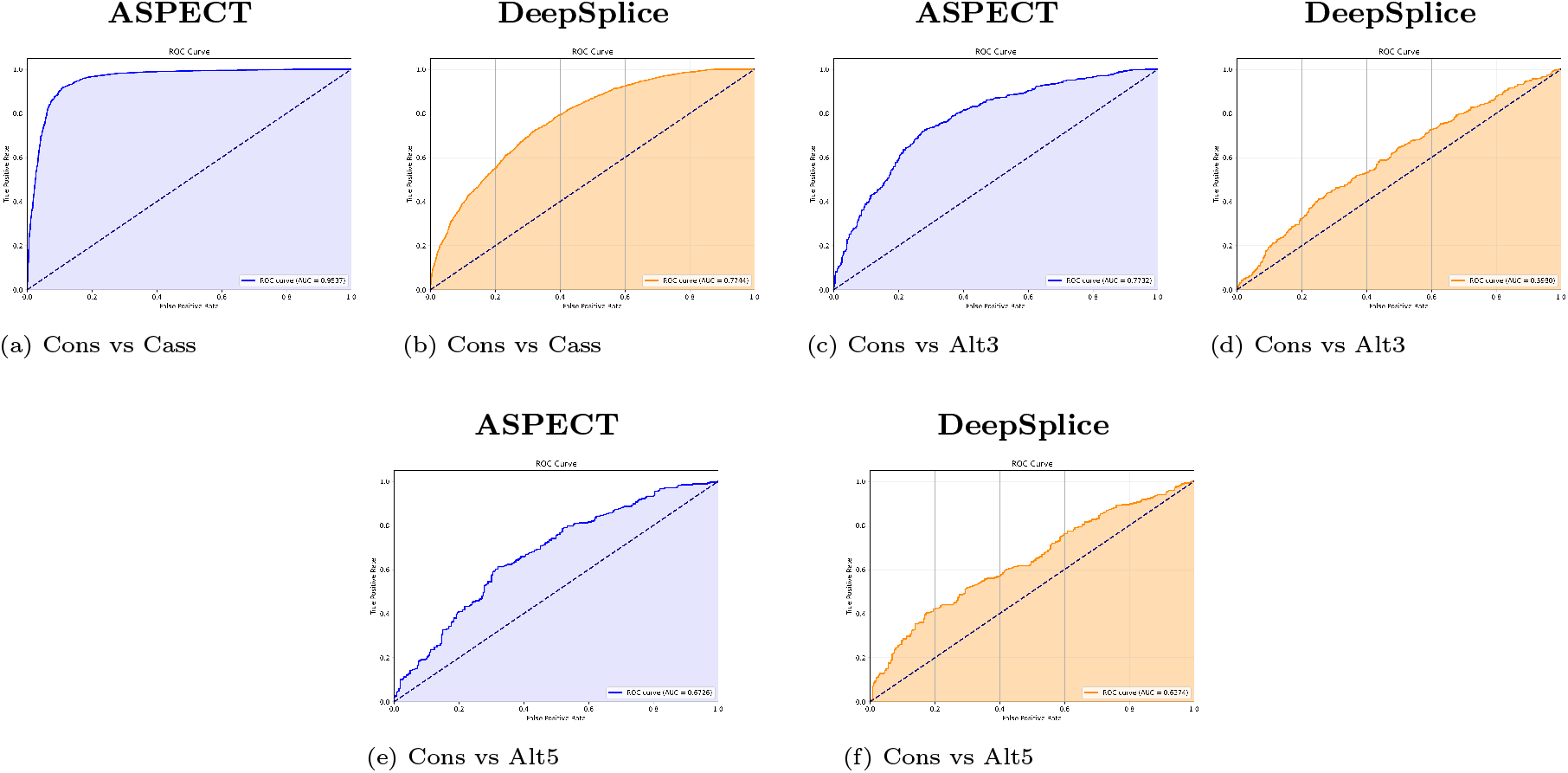
AUC curve comparisons between ASPECT and DeepSplice across three binary classification tasks: (a,b) constitutive (Cons) vs cassette (Cass), (c,d) constitutive (Cons) vs alternative three (Alt3), and (e,f) constitutive (Cons) vs alternative five (Alt5). ASPECT results are shown in blue, while DeepSplice results are shown in orange.

In addition to AUC-based evaluation, we further assess model performance using the F1-score, which provides a balanced measure of precision and recall and is particularly informative for alternative splicing event classification. Table 3 summarizes the F1-score comparison between ASPECT and DeepSplice across multiple binary alternative splicing event-pair classification tasks.

**Table 3.**
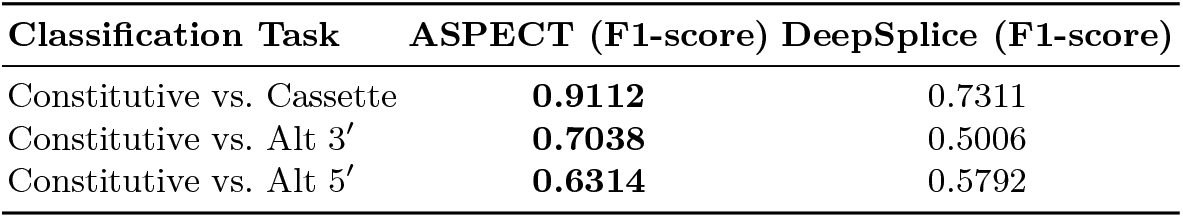
Comparison of F1-score performance between ASPECT and DeepSplice on binary splicing event pair classification tasks.

| Classification Task | ASPECT (F1-score) | DeepSplice (F1-score) |
| --- | --- | --- |
| Constitutive vs. Cassette | **0.9112** | 0.7311 |
| Constitutive vs. Alt 3' | **0.7038** | 0.5006 |
| Constitutive vs. Alt 5' | **0.6314** | 0.5792 |

Across all evaluated tasks, ASPECT achieves higher F1 Scores than DeepSplice, reflecting a better balance between sensitivity and specificity. The largest performance gains are observed for the Constitutive vs. Cassette and Constitutive vs. Alternative 3^*′*^ splice-site tasks, suggesting that ASPECT’s sequence representations more effectively capture discriminative splicing signals across a broader genomic context. These findings complement the AUC analysis and demonstrate that ASPECT provides robust classification performance relative to a state-of-the-art baseline under realistic, model-specific input configurations.

### 3.2 Hierarchical Top-*k* Inference for Alternative Splicing Event Identification

To identify dominant alternative splicing events from sequence data, we employ ASPECT within a hierarchical top-*k* inference framework (Figure 6), processing one sequence at a time. The pipeline begins with ASPECT-3, which performs three-class classification over alternative splicing event types and outputs posterior probabilities for each class. Rather than making an early hard assignment, the two most probable event labels are retained, and the sequence is routed to the corresponding ASPECT-2 model, which performs binary classification to refine the prediction between the selected event pair. This hierarchical inference strategy enables robust identification of the two most probable alternative splicing events and their relative prevalence across the analyzed dataset. Constitutive splicing events are excluded from this framework, as cancer-associated splicing dysregulation is predominantly driven by aberrant alternative splicing rather than constitutive exon usage [24, 25].

**Fig. 6:**
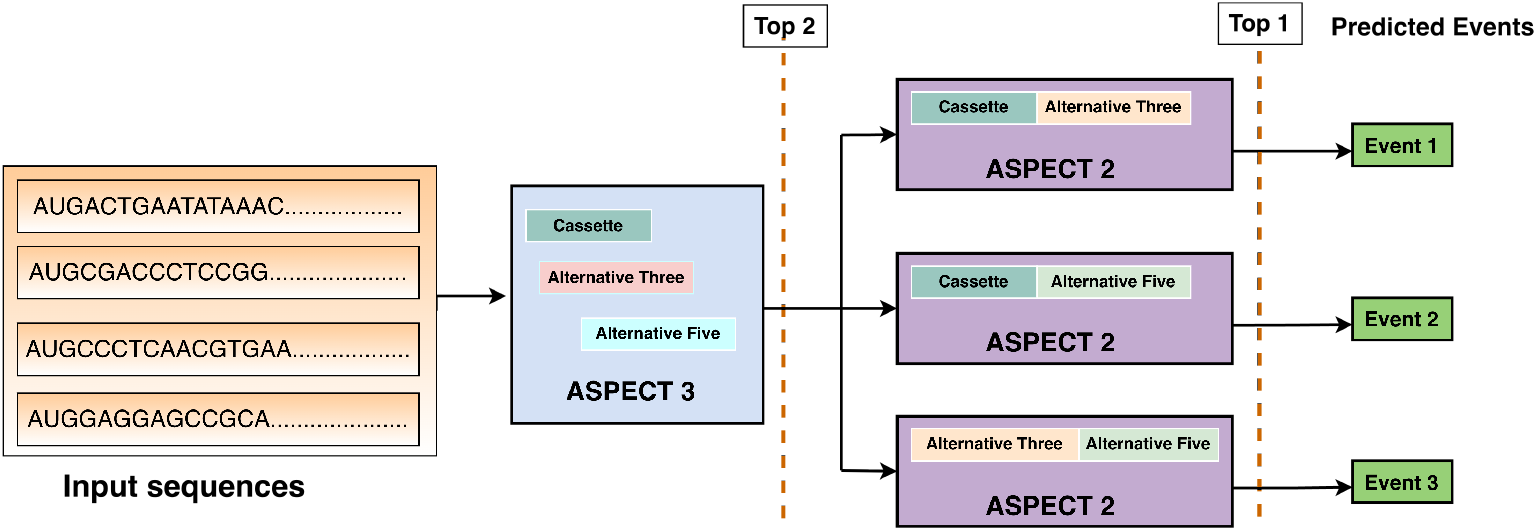
Hierarchical ASPECT transformer cascade for identifying dominant alternative splicing events from sequence data. In the first stage, ASPECT-3 performs three-class classification and assigns posterior probabilities to alternative splicing event types. For each sequence, the two most probable event labels are passed to the corresponding ASPECT-2 model, which refines the prediction through binary classification within the selected twoclass subset. Aggregating predictions across all sequences yields the two dominant splicing events and their prevalence across the dataset. Constitutive splicing events are excluded, as alternative splicing patterns are more prevalent and biologically relevant in disease and cancer contexts.

This cascaded strategy leverages the empirical observation that classification difficulty often increases as the number of output classes grows, particularly when classes exhibit subtle or overlapping patterns [26, 27]. By reducing class complexity in a stagewise manner, each model operates within a more separable decision space, improving robustness while mitigating error propagation. When applied to large collections of splicing events, this inference pipeline enables accurate estimation of event prevalence while preserving probabilistic ranking of biologically plausible splicing outcomes (Fig. 7).

**Fig. 7:**
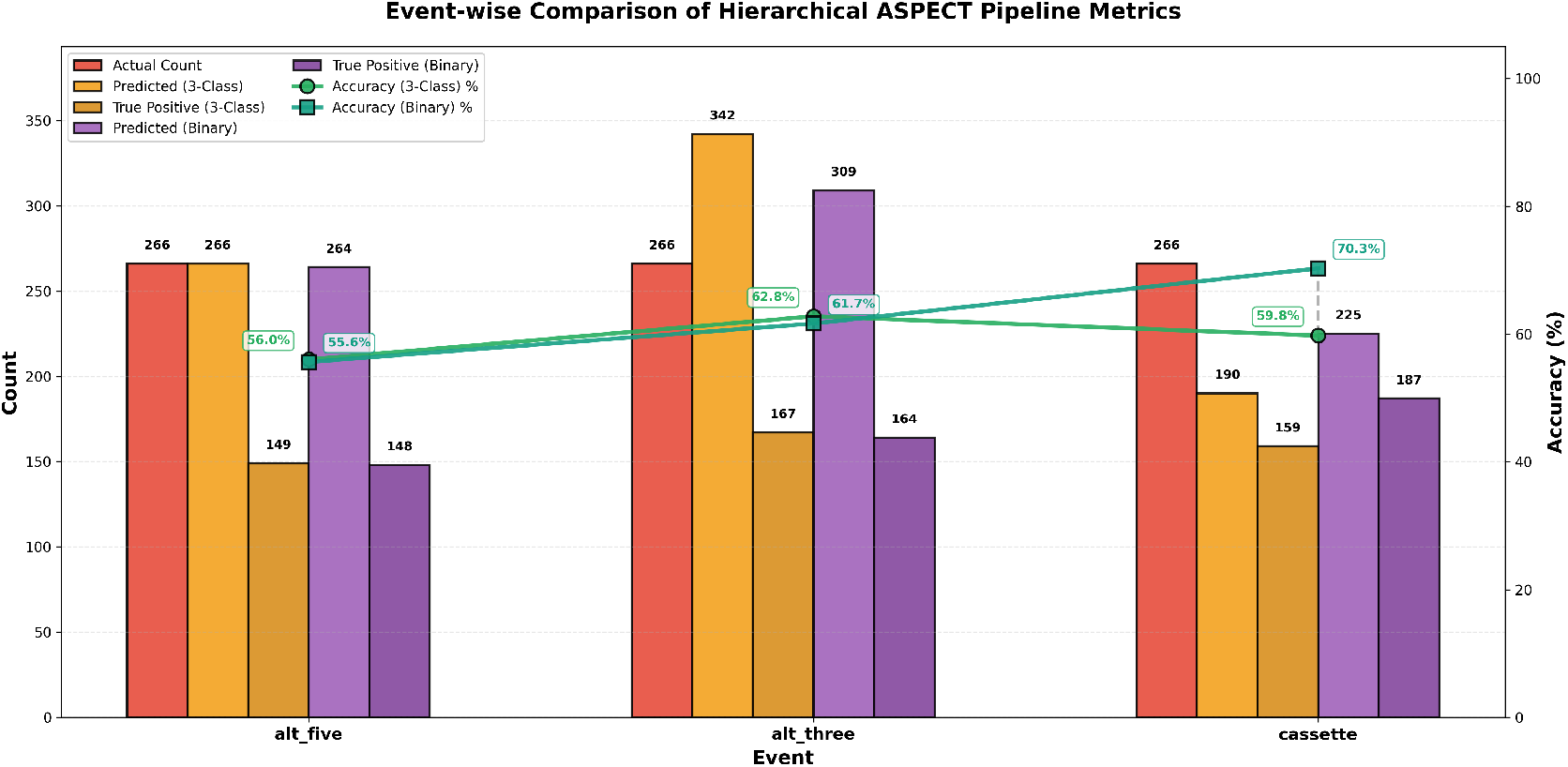
Performance metrics comparison between baseline three-class classification (59.52% accuracy) and the hierarchical ASPECT pipeline (62.53% accuracy, +3.01 pp improvement) across alternative splicing event types. Bars represent actual counts (red), predicted counts, and true positive counts with accuracy percentages. The hierarchical approach combines three-class and binary classifiers using an optimized hybrid ensemble strategy, achieving a 10.5 percentage point improvement in cassette event classification (59.8% → 70.3%).

Consistent with this design, the hierarchical ASPECT pipeline yields measurable performance gains over the baseline three-class classifier (Fig. 7). Overall classification accuracy improves from 59.52% to 62.53%, corresponding to a 3.01 percentage-point increase. Event-wise analysis indicates that these gains are not uniform across splicing types, with the most pronounced improvement observed for cassette exon events, where accuracy increases from 59.8% to 70.3%. In contrast, performance for alternative splicesite events remains largely stable, suggesting that hierarchical refinement is particularly beneficial for resolving exon-skipping patterns, which are both prevalent and challenging in cancer-associated splicing dysregulation.

### 3.3 Generalization of ASPECT to TCGA BRCA Cancer-Associated Splicing Data

To evaluate the applicability of ASPECT in a cancer-related setting, we applied the model to alternative splicing event annotations obtained from TCGA SpliceSeq [28] for breast cancer (BRCA) tumor samples [29]. TCGA SpliceSeq derives event-level alternative splicing annotations and percent spliced-in (PSI) values from RNA-seq data; however, it does not provide explicit genomic splice junction coordinates or corresponding DNA sequence context. As a result, SpliceSeq data were used exclusively for application-level evaluation and not for model training.

ASPECT was trained solely on canonical alternative splicing events derived from the HexEvent database, which are defined using genomic splice junction coordinates and sequence-based features. To enable sequence-based inference on BRCA SpliceSeq events, we reconstructed reference genome sequence representations corresponding to SpliceSeq annotated splicing events using exon coordinate information and the hg19 reference genome. This reconstruction preserves the original event definitions reported by SpliceSeq while providing fixed-length sequence inputs compatible with the ASPECT architecture. Although the training and application datasets differ in annotation origin and sequence derivation, this strategy allows direct assessment of ASPECT’s generalization to cancer-associated splicing patterns.

To generate sequence-level inputs from TCGA SpliceSeq annotations, we downloaded the BRCA splicing event matrix (`PSI_download_BRCA.txt`) from the TCGA SpliceSeq resource, which provides event identifiers, splice event types, and exon-level event definitions. Exon genomic coordinates corresponding to these events were obtained from the TCGA SpliceSeq gene structure archive (`TCGA_SpliceSeq_Gene_Structure.zip`). Using these coordinates, exon DNA sequences were extracted from the hg19 reference genome. Event-level sequences were then constructed by concatenating exon sequences according to the SpliceSeq, defined event structure, followed by length normalization to a fixed window of 1024 bp via trimming or padding. Events for which complete exon sequence information could not be recovered were excluded from downstream analysis. We note that the resulting sequences represent reference genome-based approximations of cancer-associated splicing events rather than patient-specific genomic sequences, as TCGA SpliceSeq does not provide sample-level DNA sequence information.

Figure 8 summarizes the event-wise performance of the hierarchical ASPECT inference pipeline on the BRCA SpliceSeq dataset, comparing predicted event counts and true positive detections against SpliceSeq-derived event frequencies for cassette exons, alternative 3^*′*^ splice sites, and alternative 5^*′*^ splice sites. Relative to the baseline three-class classifier, the hierarchical pipeline improves overall classification accuracy from 46.56% to 52.18%.

**Fig. 8:**
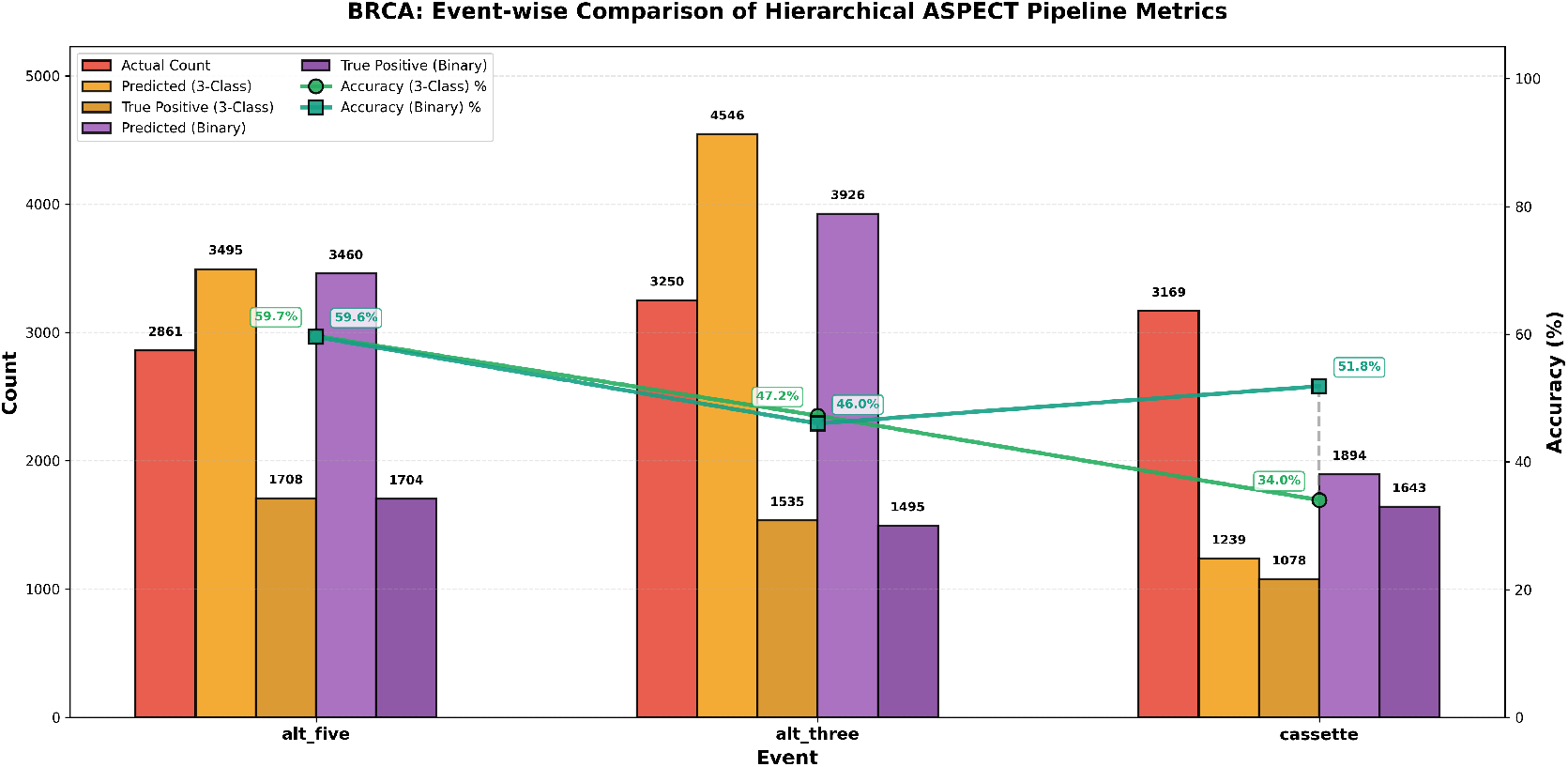
BRCA: event-wise performance comparison between the hierarchical ASPECT pipeline and the baseline three-class classifier. For each alternative splicing event type (cassette exon, alternative 3^*′*^ splice site, and alternative 5^*′*^ splice site), the bar plot compares ground-truth event counts, predicted counts, and true positive counts. On the BRCA-derived test set, the hierarchical pipeline improves overall accuracy from 46.56% to 52.18%, with cassette event accuracy increasing from 34.0% to 51.8%, while performance for alternative 3^*′*^ and alternative 5^*′*^ splice site events remains largely unchanged.

Event-level analysis indicates that these gains are driven primarily by improved identification of cassette exon events, where accuracy increases substantially from 34.0% to 51.8%. In contrast, performance for alternative 3^*′*^ and alternative 5^*′*^ splice site events remains largely stable, suggesting that hierarchical refinement is particularly effective for resolving exon-skipping patterns in cancer-derived splicing data.

Collectively, these results demonstrate that sequence representations learned from canonical splice junction annotations remain informative when applied to reconstructed cancer-associated splicing events derived from transcriptome-based annotations, supporting the robustness and transferability of ASPECT in a cancer splicing analysis context.

## 4 Conclusion

In this study, we introduced ASPECT, a transformer-based framework for alternative splicing event classification that leverages DNABERT-2 with Byte Pair Encoding (BPE) tokenization. By enabling efficient modeling of extended splice junction sequences, ASPECT captures both local splicing signals and long-range regulatory context, thereby improving discrimination across multiple alternative splicing event types. The incorporation of refined data filtering strategies and carefully selected feature thresholds further enhances classification reliability by reducing ambiguity between constitutive and highly included alternative exons.

Overall, ASPECT provides a scalable and biologically informed approach for sequence-based splicing analysis. Future work will explore tissue and condition-specific model adaptation, as well as the integration of complementary regulatory information such as epigenetic and chromatin accessibility signals, to further improve the accuracy and interpretability of splicing predictions.

## 5 Funding

This work was supported by the National Institutes of General Medical Sciences of the National Institutes of Health under award number R35GM150402 to Oluwatosin Oluwadare.

## 6 Data Availability

The open-source code and detailed documentation are available at https://github.com/OluwadareLab/ASPECT. The data and models are available at this link: https://doi.org/10.5281/zenodo.18283327

## Notes

### Competing Interest Statement

The authors have declared no competing interest.

